# Functional response predicts invasiveness but not trophic impact

**DOI:** 10.1101/2025.07.26.666957

**Authors:** Marine A. Courtois, Chloé Souques, Yann Voituron, Loïc Teulier, Vincent Médoc, François-Xavier Dechaume-Moncharmont

**Affiliations:** Universite Claude Bernard Lyon 1, CNRS, ENTPE, LEHNA, UMR 5023, Villeurbanne, F-69100, France; Université Côte d’Azur, INRAE, CNRS, ISA, Sophia Antipolis, France; Equipe de Neuro-Ethologie Sensorielle (ENES), Centre de Recherche en Neurosciences de Lyon (CRNL), CNRS, INSERM, Universite de Lyon/Saint-Etienne, Saint-Etienne, France

**Author notes:** **Corresponding author**: Marine A. Courtois. Equal contribution (Marine A. Courtois, Chloé Souques). Equal contribution (Vincent Médoc, François-Xavier Dechaume-Moncharmont). Marine A. Courtois, Chloé Souques, Yann Voituron, Loïc Teulier, Vincent Médoc, François-Xavier Dechaume-Moncharmont.

**Keywords:** Alien species, Attack rate, Biological invasions, Ecological filtering, Feeding rates, Freshwater fish, Handling time, Impact prediction, Temperature

## Abstract

Biological invasions are a major driver of biodiversity erosion mainly because invasive species show greater trophic impact than their non-invasive counterparts. The experimental paradigm for assessing this trophic impact is the functional response (FR) test that describes the relationship between *per capita* consumption rate and resource density. Two key parameters are then assessable and comparable between populations and species: the space clearance rate (attack rate, *a*) measuring predatory efficiency at low prey densities, and handling time (*h*) representing the time required to capture, handle, and digest prey.

This test is frequently conducted to compare non-invasive and invasive species and shows that invasive species have a higher FR than non-invasive species (characterized by higher space clearance rates and lower handling times) which would explain both their invasion success and their ecological impact. However, it appears that whether FR parameters differ between invasive species sampled in their native versus invasion range has never been tested, implicitly assuming that FR measures can be extrapolated to the entire range of distribution.

Using a phylogenetically corrected comparative analysis of 269 FR observations from 45 freshwater fish species (23 non-invasive species and 22 invasive species), we confirm that invasive species exhibited higher FR than non-invasive species. However, this pattern holds true only when considering invasive species sampled in their native range. Invasive species studied in their invasion range displayed functional responses comparable to non-invasive species, with similar space clearance rates and handling times. Additionally, space clearance rates decreased with temperature in non-invasive species but tended to increase in invasive species from invasive introduction ranges, suggesting that climate warming may exacerbate competitive asymmetries.

Together, these results indicate that high FR predispose species to invasiveness, but also challenge the assumption that FRs measured in the native range of a species can be directly extrapolated to predict its trophic impacts elsewhere. Our findings call for greater consideration of biogeographic context when using functional responses to assess invasion risk and ecological impact.

**Graphical abstract:** 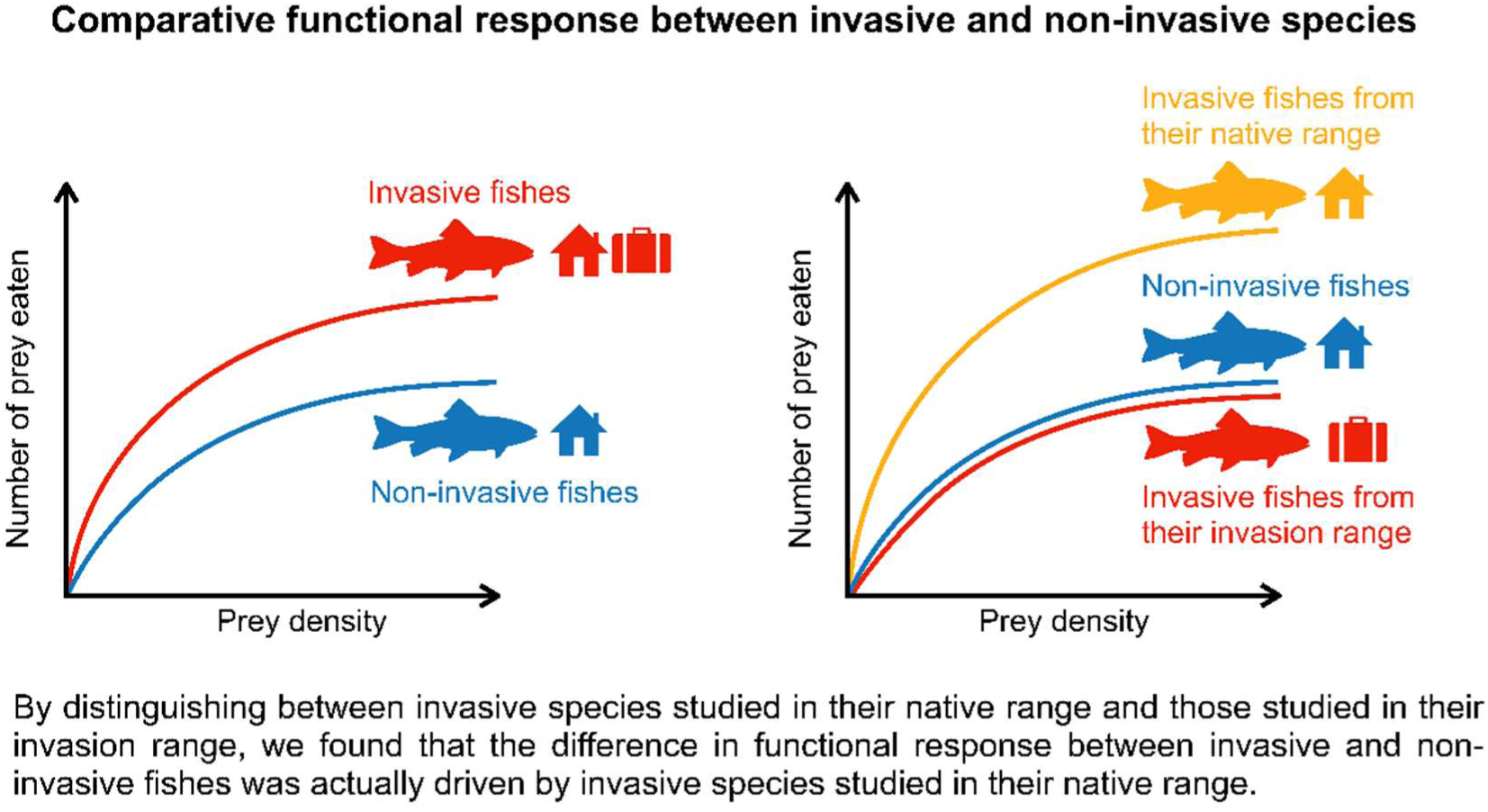

## INTRODUCTION

The interplay between species traits and the biological invasion processes continues to raise fundamental questions about ecological adaptation and species success, and it has far-reaching implications for biodiversity conservation and ecosystem resilience. Certain traits, such as selfing capacity and phenotypic plasticity in plants (Davidson et al. 2011; Razanajatovo et al. 2016), high metabolic rates (Lagos et al. 2017) or bold and aggressive behavioral types in animals (Chapple et al. 2012; Damas-Moreia et al. 2019; Jolles et al. 2020; Juette et al. 2014) can be positively selected by invasion processes, rendering the species that possess these traits potentially successful invaders. Post-establishment, species traits may be altered in response to new selective pressures within the recipient ecosystem, thereby inducing alterations in the environmental niche (niche shift, Broennimann et al. 2007; Guisan et al. 2014). For instance, the absence of natural enemies, including competitors, predators and parasites, has been identified as an important driver of invasion success (the Enemy Release Hypothesis, Colautti et al. 2004; Heger & Jeschke 2014; Meijer et al. 2016). Random variations in species traits may also be attributed to the founder effect, whereby introduced populations experience a genetic bottleneck whose magnitude is determined by population size, introduction frequency, and distance from source population (Dlugosch & Parker 2008; Dlugosch et al., 2015; Simberloff 2009). As a consequence of these non-mutually exclusive adaptive and neutral processes, species traits can diverge not only between native and invasion geographic ranges but also between introduced populations within the invasion range (Blossey et al. 1995; Björklund et al. 2010; Boets et al. 2019; Chen et al. 2023; Richards et al. 2006).

Among the traits likely to promote invasion success, those associated with resource acquisition have been extensively studied. Consumers exhibiting high attack rates (*i.e.,* effective at finding the resource when it is scarce) and/or broad diets are more likely to satisfy their nutritional needs in novel environments but also disrupt the balance of local communities. Predation by invasive species is recognized as a major threat to biodiversity and has been estimated to cause a 21% decline in species richness in aquatic habitats (Mollot et al. 2017; Reid et al. 2019). In the invasion ecology terminology, the Resource Consumption Hypothesis posits that invasive species are more efficient consumers than non-invasive species (Faria et al. 2025; Ricciardi et al. 2013). However, whether this represents a general pattern and functions as a cause or consequence of the invasion process remains to be elucidated.

A validated methodology for quantifying trophic interaction strength consists in modeling the functional response (FR), which is defined as the relationship between *per capita* consumption rate and resource density (Holling 1959). The shape of this response provides insights into potential consequences for resource populations, with certain shapes indicating more destabilizing effect than others (type-II versus type-III FR, Daugaard et al. 2019; Uszko et al. 2015). Additionally, it provides an overview on consumers’ behavior and how they attempt to maximize energy intake while minimizing costs under conditions of resource limitation (Galarowicz & Wahl 2005; Oyugi et al. 2012; Murray et al. 2013). The most prevalent FR shape is characterized by a saturation curve (Holling 1959) wherein the maximal consumption rate is constrained by handling time, which is defined as the time required for a consumer to subdue, ingest, and digest a resource item (DeLong 2021). The consumption rate at low resource densities provides an indication of consumer efficiency and is referred to as the space clearance rate or attack rate (DeLong 2021; Jeschke et al 2002; Hassell & May 1973; Holling 1959).

Under the Resource Consumption Hypothesis, invasive species are predicted to have higher FR than non-invasive species (Bollache et al. 2008; Dick et al. 2013, 2014). If invasive species are predisposed toward strong trophic impacts, populations of the invasive species in the donor area should exhibit higher FR than trophic counterparts exhibiting no invasive tendency (Dick et al. 2014). Under these circumstances, FR could be used as a screening mechanism for species invasiveness. Conversely, if trophic impact is amplified post invasion in response to new environmental conditions, such as prey naïveté (Cox & Lima, 2006; Sih et al. 2010), introduced populations should exhibit higher FR than non-invasive counterparts with which they will compete (Xu et al. 2016). From this non-exclusive alternative perspective, comparative FR is a valuable tool for quantifying the trophic impacts on local resources.

Among the growing number of FR comparisons made on invertebrates and vertebrates , several studies have reported that invasive species exhibit higher space clearance rates and lower handling times than non-invasive species, *i.e.*, trophic analogs from the introduced range without invasive status (Alexander et al. 2014; Barrios-O’Neill et al. 2014; Ens et al. 2021; Howard et al. 2018; 2022; Kattler et al. 2023; Landi et al. 2022). A recent large-scale comparative study across a broad range of taxa suggests that this may be a general pattern (Faria et al. 2025), although such studies remain relatively scarce. This scarcity is primarily due to the absence of unifying experimental framework, resulting in a divergence among FR experiments, particularly in aspects related to the characteristics of the interacting species (size, mass, density), experimental protocol (duration of the test, area, volume and shape of the experimental arena) and the environmental conditions (most notably temperature, which is of primary importance for ectotherm species). This heterogeneity posed considerable challenges to comparative FR until the development of a database that systematically collected standardized FR data from studies conducted worldwide. The FoRAGE database (Uiterwaal et al. 2018) facilitates the investigation of the role of species and environmental characteristics as potential drivers of FR parameters. Using over 300 fish FR from this database, Buba et al. (2022) found that the alien status was not a strong predictor of FR parameters. However, the majority of the FR for invasive species were obtained from populations of their native range. If, in general, invasive species do indeed have higher FR than non-invasive species, then the results of Buba et al. (2022) would suggest that this asymmetry is driven by invasive species from their (tested in) invasion range. This would support the hypothesis that increased trophic impact may emerge post-invasion as a consequence of new selective pressures in the invasion range. Nevertheless, the broader question of whether invasive and non-invasive species inherently differ in their FR remains unresolved.

The objectives of this study are twofold. Firstly, we addressed the prevailing hypothesis that invasive species exhibit higher FR than non-invasive species. Secondly, the study aimed to establish whether FR are consistent when studies are based on invasive species sampled in either their native or their invasion range. Assuming that ecological novelty (e.g., prey naïveté) exerts a more significant influence than preadaptation in increasing the trophic impact of invasive species, we predicted that, invasive species from their invasion range would demonstrate a higher FR (i.e., higher space clearance rate, lower handling time, or both) than non-invasive species and invasive species from their native range. Should such a discrepancy be observed, it would imply that results obtained from tests carried out on individuals sampled in their native range should be interpreted with caution when employed to predict the trophic impact of a species in its invasion range.We conducted this comparative analysis in 45 freshwater fish species including 23 non-invasive species and 22 invasive species, with invasive species represented by populations sampled from either their native or their invasion ranges. Freshwater fish species are particularly sensitive to biological invasions and alterations to their trophic networks (Albert et al. 2021; Reid et al. 2019; Sayer et al. 2025). They are among the species for which the most data on FR are available (Uiterwaal et al. 2018). The analysis of phylogenetic inertia was facilitated by the focus on a single, homogeneous taxonomic group.

## MATERIAL AND METHOD

### Species Selection and Classification

We conducted a comparative analysis by cross-referencing two databases (See PRISMA flowchart, Fig. S1). First, we used the ‘FoRAGE’ database (see details below) to identify all freshwater fish species for which FR parameters had been measured. We then used the ‘GRIIS’ (see details below) database to determine the invasive status of each species.

### Functional response data

FR parameters used in this analysis were obtained from the FoRAGE database (version 5 on 20 December 2024, Uiterwaal et al. 2018, Uiterwaal et al. 2022), which is an extensive compilation of FR derived from extant literature. These FR parameters have been standardized to ensure the comparability of the space and time units (see details below). The median values were reported alongside the 95% quantiles of the bootstrapped parameter distributions. The database, when available, also reported information regarding predators (species, body mass, life stage, juvenile or adult), prey (species, body mass, life stage), and the original experimental conditions (temperature, arena dimension). A total of 353 observations of FR in freshwater fish species were identified in FoRAGE database (See PRISMA flowchart, Fig. S1). Their FR were fitted in the database with a type II model using standardized methods (Uiterwaal et al. 2018). The type II FR is described by Holling’s disk equation (Eq. 1, Holling 1959):

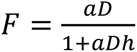

This FR equation relates consumption rate F (number of preys consumed per unit of time) to prey density D (number of preys per unit of space, where space is generally a volume of water in the case of fish species). FR is determined by two key parameters, *a* and *h*. The space clearance rate or attack rate *a* (DeLong 2021; Holling 1959) defines the volume of space cleared of prey per predator. It serves as a measure of predatory efficiency at low prey densities (Jeschke et al. 2002). The higher the value of *a*, the steeper initial slope for the functional response curve. The handling time *h* corresponds to the time required for a predator to capture, handle, and digest a prey item (DeLong 2021). The smaller the value of *h*, the greater the height of the plateau on the functional response curve.

In the majority of FR studies, the units of the parameters *a* and *h* remain undefined. Authors typically compared responses of several populations within the same species and the same study using identical experimental setup. As the primary focus of these studies was on the significance of the difference between homogeneous experimental groups, direct comparisons were made between the numerical values of the estimates, albeit without a clear definition of the units. Nevertheless, it is imperative to unambiguously define these units in order to allow meaningful comparisons across studies. In order to express the FR in homogeneous units, the focus was placed on studies based on 3D arena, i.e. where space units were volumes (See PRISMA flowchart, Fig. S1).

In accordance with the standardized value reported in FoRAGE database, the space clearance rate *a* was expressed in cubic meters of water per day per predator (units: m^3^.day^−1^). The handling time *h* was expressed in time unit times number of predator per number of prey (units: day). Our primary analyses were conducted using these per capita units (hereafter referred to as ‘standard units’), which focused on the number of prey individuals removed from the environment. However, the number of prey items consumed varied by several orders of magnitude, depending on whether the prey species in question are, for instance, *Daphnia* or vertebrate species. From the perspectives of functional ecology or predator physiology, the consequences of consuming one prey item are not equivalent when the predator and prey species strongly differ in size. To account for substantial variability in body size, analyses were repeated after standardizing parameters *a* and *h* by prey and predator biomass (hereafter referred to as ‘biomass units’). In these secondary analyses, biomass clearance rate *a* was expressed in cubic meters per kilogram of predator per day (units: m^3^.kg^−1^.day^−1^), and biomass handling time *h* was expressed in time unit times kilogram of predator per kilogram of prey per day (units: day).

The analyses presented herein were based on a total of 269 estimates of a and h, reported in a total of 45 freshwater fish species from 40 studies (See PRISMA flowchart, Fig. S1). Interspecific differences were analyzed through a comparison of their mean values of the FR parameters *a*, or *h*, but also the FR Ratio FRR = *a/h*, which has recently emerged as a reliable metric for balancing the two FR parameters (Faria et al. 2025; Cuthbert et al. 2019b). Ecologically, high trophic impacts are predicted for species exhibiting high functional responses, whether driven by high space clearance rates (*a*), low handling times (*h*), or high FRR. Due to missing information about prey body mass in the original articles, the analyses based on biomass units were conducted on 254 estimates for the handling time *h* and FRR *a/h*.

### Invasive status and geographic range of sampling

The FR database was completed with the invasive status of each freshwater fish species specified as ‘invasive’ (IS) or ‘non-invasive’ (NIS). For this purpose, we relied on the Global Register of Introduced and Invasive Species (GRIIS, Pagad et al. 2018), conceived within the framework of the Global Invasive Alien Species Information (GIASI) partnership of the Convention on Biological Diversity (CBD). It provides the first country-by-country checklists of invasive species. This classification has been established on the basis of the number of countries in which the species have been introduced and their ecological impact in the countries into which they were introduced. The classification of a species as ‘invasive’ was determined by its introduction into at least one country outside the limits of its native geographic range, and by the potential threat to native biodiversity that its spread poses, which is referred to as “evidence of impact” in GRIIS database. Following this classification, our study was based on a total of 22 invasive species from 31 studies (168 observations FR estimates for *a* and *h*), and 23 non-invasive species from 19 studies (101 observation of *a* and *h*). Furthermore, we distinguished between the geographical regions (’native range’ vs invasion range’) from which fish were sampled. All non-invasive species (NIS) were sampled in their native range. However, for invasive species (IS), it was necessary to differentiate between studies conducted on invasive species sampled in their native range (IS-NR, donor population, 13 species from 20 studies, 107 estimates) and those conducted on invasive species sampled in their invasion range (IS-IR, introduced population, 11 species from 13 studies, 61 estimates). The invasive status (NIS, IS-NR or IS-IR) of each FR observation was established by comparing the sampling geographic range reported in each original study with the corresponding invasive status and native range referenced in the GRIIS database. All observations identified in the FoRAGE database that did not report information regarding the sampling geographic range of the fish species have been excluded from the analyses (see PRISMA flowchart, Fig. S1). Only three invasive species (*Gambusia affinis*, *Lepomis macrochirus*, and *Poecilia reticulata*) have been studied in both their native and invasion ranges.

### Effect of the temperature

A previous study (Buba et al. 2022) showed an effect of the temperature at which the FR assays were carried out, so temperature was included in our analyses. Observations from the FoRAGE database lacking experimental temperature information were excluded (see PRISMA flowchart, Fig. S1).

### Phylogenetic tree

Species that share a common ancestor often possess inherited traits due to descent rather than independent adaptation. Some traits exhibit a strong phylogenetic signal, meaning they are conserved along lineages. Consequently, closely related species cannot be considered as independent data points. Additionally, certain taxa may be overrepresented in the literature, potentially biasing analyses. Comparative analyses must incorporate phylogeny to enhance robustness and reduce biases (Cinar 2022; Lajeunesse et al. 2013). Our analyses accounted for the phylogenetic relationships between the fish species by integrating a species-level phylogenetic correlation matrix in all models (see statistical analysis section for details). This matrix was derived from the phylogenetic tree (Fig. 1) of the 45 freshwater fish species present in our dataset, with the R package ‘FishPhyloMaker’ (Nakamura et al. 2021). Invasive species were distributed across all families or orders, with no taxon disproportionately represented (Fig. 1). Lynch’s formula estimated the phylogenetic signal (or phylogenetic heritability) for each FR parameter expressed in both standard and biomass units (Garamszegi 2014).

**Figure 1:**
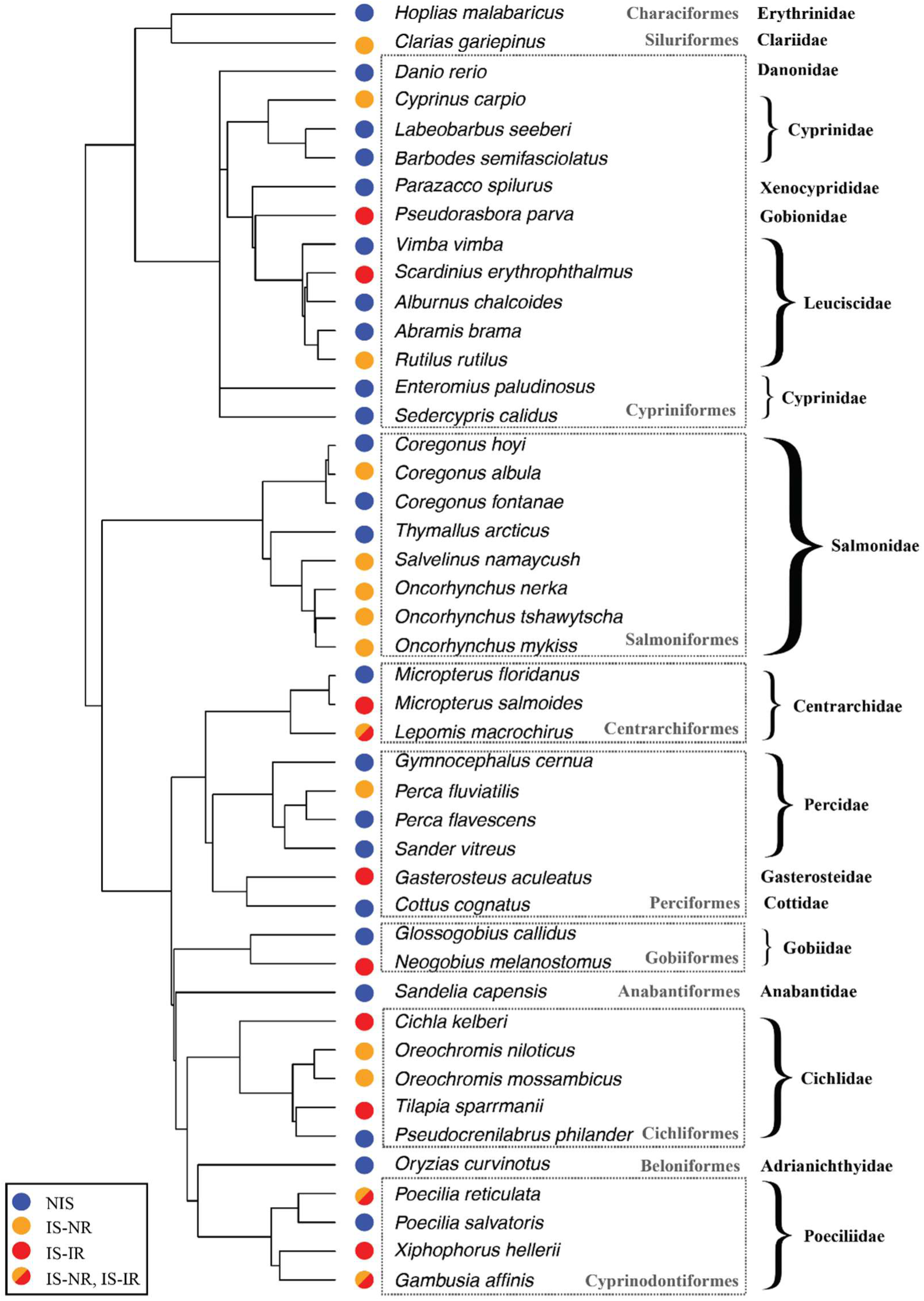
Phylogenetic tree of the 45 freshwater fish species used in the analyses. Each colored circle in front of the species names illustrates the status of the species: non-invasive species (NIS, blue circle, all were sampled in their native range), invasive species sampled either in their native range (IS-NR, donor populations, orange circle) or in their invasion range (IS-IR, introduced populations, red circle). Some invasive species have been sampled both in their native range and in their invasion range depending on the specific study undertaken (IS-NR and IS-IR, bicolor circle). Taxonomic rank is reported at the order (in grey in dotted boxes) or family level (in bold black, on the right side).

### Ethical approval

This study used exclusively publicly available data from previously published sources. No new data were collected, and no animals were handled or manipulated by the authors. Therefore, ethical approval was not required.

### Statistical analysis

All analyses were carried out separately for each FR parameters (*a*, *h*, FRR, both in the standard units and in biomass units). Due to the wide variation between species, the distributions of these parameters were highly skewed. We used log transformation to normalize data and make it symmetric, thus meeting the assumptions of normality and homoscedasticity. There were no missing data in our datasets, thereby enabling model selection based on information theory (Nakagawa & Freckleton 2011) to test whether invasive status (non-invasive vs. invasive species) or sampling status (NIS, IS-NR, IS-IR) explained the differences in FR parameters.

The phylogenetic correlation matrix derived from our phylogenetic tree was used as random variable in all models using a phylogenetic mixed model approach (Hadfield & Nakagawa 2010) with the R package MCMCglmm for Bayesian mixed models (Hadfield 2010), assuming a Brownian model of trait evolution (Hadfield & Nakagawa 2010). The Markov-chain Monte Carlo (MCMC) was used with 3×10^5^ iterations, 300 burn-in period and a thinning interval of 100 and an inverse-Wishar prior. Prior type had no impact on the conclusions that were drawn from the analyses.

For each FR parameter (*a*, *h*, or FRR) and for each unit (standard or biomass unit), several competitive models were considered. The simplest model, ‘Model 0’, solely took into account phylogenetic correlation matrix as random variable. No discrimination was made between invasive status (non-invasive vs. invasive species) or sampling status (NIS, IS-NR, IS-IR). ‘Model 1’ considered the invasive status, invasive (IS) versus non-invasive (NIS) species, as fixed effect, irrespective of the geographic range in which the species had been sampled. ‘Model 2’ considered the sampling status as fixed effect. This composite variable was a three-level factor: non-invasive species (NIS, which were always sampled in their native range), invasive species sampled in their native range (IS-NR, donor population), and invasive species sampled in their invasion range (IS-IR, introduced population). Because Model 1 and Model 2 were not nested, it was not possible to directly compare them using likelihood ratio test. They were compared using information criterion (Burnham & Anderson, 2004). Model selection was performed using the deviance information criterion (DIC, Spiegelhalter et al. 2002). The models with the lowest DIC value were considered the best models. Models with ΔDIC > 2 were considered to have substantially poorer fit, and models with ΔDIC < 2 were considered to have equivalent support to the best model (Barnett et al. 2010). All these analyses were repeated separately for each FR parameter (*a*, *h*, FRR) both in standard units and in biomass units.

The temperature was taken into account as fixed variable. ‘Model 3’ considered the temperature, the sampling status, and their interaction as fixed effects. The effects of the explanatory variables and their interaction term were analyzed by comparing the DIC of the nested models, using the same model selection procedure as described above.

Data processing and analysis were carried out with the R programming language (version 4.4.3, R Development Core Team 2025).

## RESULTS

### Effect of the invasive status

FR parameters expressed in standard units varied significantly according to the invasive status (non-invasive vs. invasive species). Invasive species had a significantly higher FR than non-invasive species, with a higher space clearance rate *a* (posterior mean = -1.07, 95% credible interval 95%CrI = [-1.63; -0.55], pMCMC = 6.7×10^−4^, Fig. 2A), a lower handling time *h* (posterior mean = 0.82, 95%CrI = [0.18; 1.54], pMCMC = 0.022), and a higher FRR = *a/h* (posterior mean = -1.8, 95%CrI = [-2.83; -0.94], pMCMC = 3.0×10^−4^). Similar effects, though less pronounced, were found for FR parameters expressed in biomass units: space clearance rate *a* (posterior mean = -0.49, 95%CrI = [-0.94; -0.028], pMCMC = 0.043), handling time *h* (posterior mean = 0.60, lower and upper 95%CrI= [-0.010; 1.23], pMCMC = 0.056), FRR a/h (posterior mean = -1.12, 95%CrI = [-2.01; -0.26], pMCMC = 0.019).

**Figure 2:**
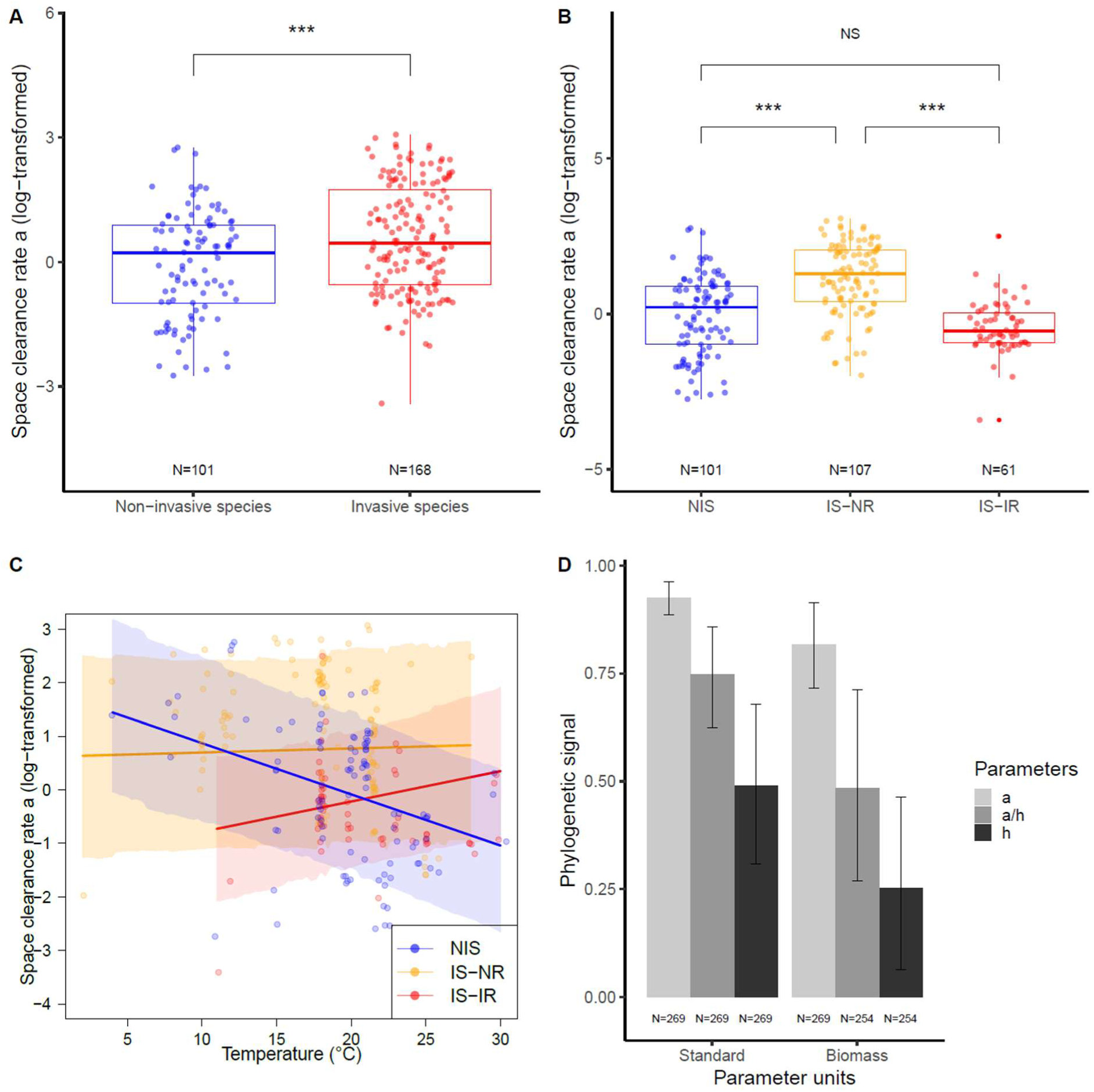
The functional response parameters are expressed in standard units (log-transformed). **A)** Boxplot of space clearance rate as a function of the invasive status, non-invasive vs. invasive species. **B)** Boxplot of space clearance rate as a function of the sampling status, which was either non-invasive (NIS, in blue, all were sampled in their native range), invasive species sampled in their native range (IS-NR, donor populations, in orange) or invasive species sampled in their invasion range (IS-IR, introduced populations, in red). **C)** Interaction between sampling status and temperature. **D)** Barplot of the phylogenetic signal for the various functional response parameters (space clearance rate a, handling time h and functional response ratio FRR = a/h) expressed in standard or biomass units. These signals were calculated using Lynch’s phylogenetic heritability formula (Garamszegi, 2014). The sample size (N) is indicated below each group when relevant. The sample size differed for the analyses in biomass unit due to unreported values. Each data point corresponds to the report in a specific article of one estimated value of space clearance rate for one species. Multiple data points may correspond to the same species studied across different articles. The mixed-effect model incorporates multiple imputations and accounts for phylogenetic inertia. *** indicates pMCMC < 0.001, and NS pMCMC > 0.05.

### Effect of the sampling status

We then categorized the invasive species according to their sampling range, either donor population (IS-NR) or introduced population (IS-IR). The ΔDIC > 6 in the model selection procedure (Table 1) indicated strong evidence that the ‘Model 2’ based on sampling status (NIS, IS-NR, IS-IR) was a substantially better fit of the data compared to ‘Model 1’ based solely on the invasive status (non-invasive vs. invasive species) for all FR parameters expressed in standard units. Donor populations had a significantly higher FR than introduced populations, with a higher space clearance rate *a* (posterior mean = 0.88, 95%CrI = [0.49; 1.27], pMCMC < 3×10^−4^ ; Fig. 2B), a lower handling time *h* (posterior mean = -1.18, 95%CrI = [-1.89; -0.44], pMCMC = 1.3×10^−3^ ; Fig. S2A) and higher FRR *a/h* (posterior mean = 2.11, 95%CrI = [1.22; 2.97], pMCMC < 3×10^−4^, Fig. S2C). Furthermore, introduced populations and native species did not differ in space clearance rate *a* (posterior mean = -0.56, 95%CrI = [-1.15; -0.02], pMCMC = 0.062, Fig. 2B), handling time *h* (posterior mean = 0.038, 95%CrI = [-0.83; 0.82], pMCMC = 0.92, Fig. S2A) and FRR *a/h* (posterior mean = -0.51, lower and upper 95%CrI = [-1.55; 0.50], pMCMC = 0.33, Fig. S2C). Fully consistent results were observed for the three parameters *a*, *h*, and FRR expressed in biomass units (Table S1, Fig. S3A,C,E).

**Table 1.** Model selection table for the influence of sampling status on each functional response parameter (space clearance rate a, handling time h, and functional response ratio FRR = a/h) expressed in standard units. The competitive models were ranked according to their DIC. For a detailed description of the three models, refer to the main text.

| Model | Space clearance rate $a$ | | Handling time $h$ | | FRR = $a/h$ | |
| --- | --- | --- | --- | --- | --- | --- |
| | DIC | ΔDIC | DIC | ΔDIC | DIC | ΔDIC |
| Model 2 (sampling status) | 579.48 | 0 | 1052.58 | 0 | 1105.65 | 0 |
| Model 1 (invasive status) | 597.94 | 18.46 | 1058.67 | 6.09 | 1121.22 | 15.57 |
| Model 0 (intercept only) | 598.97 | 19.49 | 1059.72 | 7.14 | 1124.03 | 18.38 |

We accounted for phylogenetic non-idenpendance in our analyses. However, although the family Salmonidae was well represented in our dataset, no FR observations were available for invasive Salmonidae in their invasion range (IS-IR, Fig. 1). To ensure that the presence of this taxon did not bias our conclusions, we repeated the previous analyses on a reduced dataset excluding the Salmonidae. The results from these complementary analyses were fully consistent with those obtained from the complete dataset (see Fig. S4 in standard units and Fig. S5 in biomass units).

For three species, FR were quantified in both native and invasion ranges, enabling a paired species-level comparison. Linear models (log-transformed *a*, *h*, and *a/h*) were fitted separately for each species. For *Lepomis macrochirus* and *Poecilia reticulata*, the results were consistent with the results observed in the overall analysis with higher FR in the donor population (IS-NR) than in the introduced one (IS-IR). For *Gambusia affinis*, the pattern could not be confirmed due to the very small number of observations (only one observation for IS-IR). Species-specific results are presented in Fig. S6 (standard units) and Fig S7 (biomass units). In the global model including the three species, with species identity as a random effect, and invasive populations separated into donor (IS-NR) and introduced (IS-IR) populations, donor populations showed higher space clearance rate *a* (posterior mean = 0.46, 95% CrI = [0.30; 0.63], pMCMC < 3×10⁻⁴; Fig. S4D), lower handling time *h* (posterior mean = 1.71, 95% CrI = [1.15; 2.30], pMCMC < 3×10⁻⁴; Fig. S4H), and higher FRR *a/h* (posterior mean = 2.06, 95% CrI = [1.38; 2.78], pMCMC < 3×10⁻⁴; Fig. S4L) than introduced populations (Fig. S6D,H,L for standard units, or Fig. S7D,H,L for biomass units).

### Effect of the temperature

For the study of space clearance rate *a*, there was substantial support for an effect of the interaction between the sampling status (NIS, IS-NR, IS-IR) and temperature (ΔDIC > 12 in Table 2). In non-invasive species (NIS), the space clearance rate *a* decreased with the temperature (posterior mean slope = -0.095, 95%CrI = [-0.136; -0.0526], pMCMC < 3×10^−4^, Fig. 2C), whereas introduced populations did not react significantly, though a non-significant positive tendency was observed (Table S3). The same pattern was observed for the FR ratio *a/h* (ΔDIC > 2 in Table 2, Fig. S2D, Table S3). By contrast, the handling time *h* (Table 2, Fig. S2B, Table S3) was unaffected by the temperature: the model including the interaction term between sampling status and temperature corresponds to ΔDIC > 3 (Table 2). Among the best models (ΔDIC < 2), the most parsimonious model does not include the temperature variable. The analyses based on the dataset in biomass units led to consistent results (Table S2, Fig. S3B,D,F, Table S4).

**Table 2.** Model selection table for the effect of sampling status and experimental temperature on each functional response parameter (space clearance rate a, handling time h, and functional response ratio FRR = a/h) expressed in standard units. The full model (‘Model 3’, see main text) and all the nested models were ranked according to their DIC. The fixed variables considered were the experimental temperature (T°), the sampling status (NIS, IS-NR, IS-IR), and their interaction term (T°:status) between temperature and sampling status. Degree of freedom (df) for each model is also reported.

| Parameter | Nested model | | | df | DIC | ΔDIC |
| --- | --- | --- | --- | --- | --- | --- |
| $a$ | $T^\circ$ | Status | $T^\circ$: Status | 8 | 565.9 | 0 |
| | $T^\circ$ | Status | | 6 | 578.8 | 12.89 |
|  |  | Status |  | 5 | 579.3 | 13.39 |
|  |  |  |  | 3 | 599.0 | 33.05 |
| | $T^\circ$ | | | 4 | 601.0 | 35.01 |
| $h$ | | Status | | 5 | 1052.6 | 0 |
| | $T^\circ$ | Status | | 6 | 1054.2 | 1.62 |
| | $T^\circ$ | Status | $T^\circ$: Status | 8 | 1055.7 | 3.11 |
|  |  |  |  | 3 | 1059.8 | 7.18 |
| | $T^\circ$ | | | 4 | 1061.7 | 9.04 |
| $FRR = a/h$ | $T^\circ$ | Status | $T^\circ$:Status | 8 | 1103.4 | 0 |
|  |  | Status |  | 5 | 1105.6 | 2.18 |
| | $T^\circ$ | Status | | 6 | 1107.8 | 4.38 |
|  |  |  |  | 3 | 1124.0 | 20.64 |
| | $T^\circ$ | | | 4 | 1126.6 | 23.17 |

### Phylogenetic signals

The phylogenetic signals were greater than 0.50 for all FR parameters, with the strongest effect observed for the space clearance rate (a = 0.93, 95%CI = [0.89; 0.96] in standard units, a = 0.82, 95%CI = [0.72; 0.91] in biomass units, Fig. 2D). A phylogenetic signal of 0 indicates phylogenetic independence, whereas a signal of 1 indicates very strong variation in the parameter with phylogeny (Freckleton et al. 2002). These results underscore the importance of including phylogeny in comparative analysis of FR in order to avoid phylogenetic bias.

## DISCUSSION

We compiled 269 FR from 45 freshwater fish species to test whether invasive species are stronger consumers than non-invasive species while accounting for the geographic range in which the invasive species were sampled, their phylogenetic relationships, and temperature. Overall, invasive species exhibited higher FR compared to non-invasive species, but surprisingly this difference was driven by the invasive species tested in their native range. These results question the paradigm that drives the comparison of FR between invasive and non-invasive species and suggest new hypotheses regarding how the invasion process can make species traits evolve.

The assessment of general patterns from literature review may be subject to publication bias (Møller and Jennions 2001, Rothstein et al. 2005, Nakagawa et al. 2021). Positive results, particularly those confirming that invasive species have superior functional responses, are more likely to be published than null or contradictory findings, representing a well-documented bias in the ecological literature. Our analysis relies on parameters from the FoRAGE database, the most extensive compilation of functional response data currently available. While the database provides standardized data across a wide taxonomic scope, it inherently reflects the publication biases present in the underlying literature. Yet, the novelty of our study lies in distinguishing invasive species from their native versus introduced ranges. Because this distinction has not been previously addressed, any publication bias would likely affect both categories equally and would not qualitatively alter our findings. Nonetheless, we acknowledge that publication bias cannot be entirely eliminated in studies based on published data, and this represents a limitation of any meta-analysis or comparative study. If anything, our results may represent an upper bound on the true effect size. When invasive species were not differentiated based on whether they were sampled in their native (donor population, IS-NR) or invasion range (introduced population, IS-IR), it was found that they had higher FR than non-invasive species, with significantly higher space clearance rates and a tendency for lower handling times. The results remained valid when the FR were expressed in ‘standard units’ (*per capita*) or in ‘biomass units’, which accounted for the wide variability in body mass of prey and predator species across studies. This is consistent with the prediction and underlying hypothesis of the comparative FR approach, which states that invasive species are better consumers than non-invasive species. This explains their success in becoming invasive and their impact on the invaded ecosystems (see the Resource Consumption Hypothesis, Faria et al. 2025; Dick et al. 2014; Ricciardi et al. 2013). Until the present work, this hypothesis had been supported by empirical FR studies in various taxa (Alexander et al. 2014; Barrios-O’Neill et al. 2014; Crookes et al. 2019; Cuthbert et al. 2019a; Howard et al. 2018 ; Laverty et al. 2015; Paterson et al. 2015; Mofu et al. 2019; Madzivanzira et al. 2021) but validation from more than one large-scale comparative approach was needed (Faria et al. 2025). However, when the geographic range in which the FR of invasive species were sampled was taken into account, it appeared that the general pattern of higher consumption rate in invasive species compared to non-invasive species was largely driven by studies based on donor populations (IS-NR).

Invasive species sampled in their native range (donor populations, IS-NR) exhibited significantly higher FR than the non-invasive species, supporting the pre-invasion scenario in which some species would be predisposed to be invasive (Cadotte et al. 2018; Hulcr et al. 2017; Schlaepfer et al. 2010; Suarez & Tsutsui 2008). In the previous and unique comparative study also using the FoRAGE database, Buba et al. (2022) combined 309 FR of freshwater and marine predatory fishes to test the influence of size, activity level, temperature and invasive status as potential predictors of FR parameters. Although species were not classified according to their sampling range (*i.e.*, native or invasion range), most invasive taxa were tested in their native range, enabling direct comparison with our study. Their results were qualitatively consistent with ours but revealed only a weak link between invasive status and space clearance rate, possibly because phylogenetic inertia was not considered.

Our result suggests that having a high FR ‘at home’ might offer a better chance of establishing a population in the invasion range, for instance by withstanding competition. However, by definition, our dataset only included species that successfully established invasive populations and formally testing this hypothesis would require FR data on failed invaders, which is unfortunately not available.

The high FR of donor populations (IS-NR) suggests that comparative FR could be used to screen species for invasiveness with the aim of identifying likely future invaders and preventing invasion (Hulcr et al. 2017; Van Kleunen et al. 2010). Theory can help explain why species predisposed to invasiveness would not cause imbalance to their community and remain unnoticed. When interspecific competition is greater than intraspecific competition, superior competitors are predicted to exclude inferior competitors (competitive exclusion, Bøhn et al. 2008; Gause 1934; Hardin 1960; Paine 1966; Tilman 1994). According to optimal foraging theory, the most successful and therefore most frequently encountered species should be the preferred resources of higher trophic-level natural enemies such as predators and parasites (Ishii & Shimada 2012; Lacher et al. 1982; Pyke 1984). As a result, natural enemies promote coexistence at the lower trophic level by decreasing the strength of competitive exclusion (Chase et al. 2002; Chesson & Kuang 2008; Stump & Chesson 2017). Competitors would exhibit their high potential once released from their natural enemies, as occurs in the invasion range or in the experimental setting of FR derivation.

Contrary to our prediction that prey novelty in the invasion range would benefit introduced populations (IS-IR) through increased predation rates, their FR ratio was, on average, equivalent to that of non-invasive species (NIS) and lower than that of donor population (IS-NR). This decrease was mainly driven by smaller space clearance rates and could result from environmental constraints operating in the recipient ecosystem and leading to trait convergence between introduced populations (IS-IR) and resident non-invasive species (Craven et al. 2018). Comparative FR studies involving donor (IS-NR) and introduced (IS-IR) populations of the same invasive species, while also accounting for the time elapsed since introduction, could help identify which mechanisms could explain trait convergence in the recipient ecosystems. Environmental filtering (Gaston et al. 2009; Keddy 1992), intra- and intergenerational phenotypic plasticity (within or between individuals), or genetic changes (Marzluff 2012) could all be relevant factors. Whether invasive species actually perform better in their invasion range remains controversial depending on the taxa studied (Parker et al. 2013). Other phenotypic traits beyond trophic impact may be influenced by ecological filters and could diverge between invasive species originating from their donor or recipient range. Indeed, if resource acquisition is primarily driven by underlying metabolic demands and covariates with particular behavioural traits (*i.e.* proactive) as outlined within unifying theories and some empirical evidence (Careau et Garland 2012; Nannini et al., 2012 ; Réale et al. 2010; Ronce et Clobert 2012; Taylor and Dunn 2018), it would be valuable to question a potential variation in invasion syndromes relative to individuals’ sampling ranges. Furthermore, the study of invasion gradients demonstrated that invasive species, whether sampled in their core or front areas, can exhibit different personality and physiological traits (Alves et al. 2025 ; Galli et al. 2023 ; Myles-Gonzales et al. 2015), suggesting that ecological filters associated with the successful establishment in the local environmental conditions may play a significant role in shaping the phenotype of invasive species. Consequently, our study underscores the necessity of adopting an integrative approach to elucidate the contribution of ecological filtering to the phenotypic plasticity observed in invasive species.

Only three invasive species have been studied in both their native (IS-NR) and invasion (IS-IR) ranges, meaning that a direct assessment of the effect of sampling status was possible for only a limited subset. These analyses were consistent with the overall analyses of the full dataset, which lends support to the robustness of our conclusions. Yet, data are currently lacking to perform such Comparative FR approaches (CFR, Cuthbert et al. 2019b; Faria et al. 2023, 2025). Expanding the number of studies that quantify FR in invasive species across both their native and invasion ranges is therefore a critical priority. Such work would enable more powerful paired analyses, reduce uncertainty associated with between-species heterogeneity, and provide a stronger empirical basis for disentangling the roles of biogeographic context and sampling status in shaping observed FR patterns.

More generally, a species may exhibit relatively low performance in its native range yet exert strong impacts in its introduced range if resident species are even less effective predators. FR should therefore be interpreted as comparative rather than absolute measures. The potential impact of an invasive species cannot be inferred solely from its absolute scores of FR, particularly when it was sampled in its native range. Instead, correct interpretation of functional responses requires comparing the functional responses of invasive species with those of non-invasive species sampled within the same geographic range.

Equivalent FR between introduced populations (IS-IR) and resident non-invasive species also challenges the general assumption of a positive relationship between resource intake and performance (Brown et al. 2004; DeLong et al. 2021 ; Kooijman 2010), in that metabolically efficient species might not be those with high feeding rates, and therefore high FR, but those able to maintain viable metabolic rates while reducing the amount of resources required. Such “thrifty” species would have more chances to persist in ecosystems where ecological niches are occupied and become integrated into the local trophic network. Despite the literature on the crucial role of physiology in shaping the evolutionary trajectories and biogeographic distributions of non-invasive species (Somero 2002, 2005), there are to date few studies aimed at assessing the physiology of invasive species. Compared to non-invasive species, they would be more eurytherms (Kelley 2014, but see Boher et al. 2018), have higher mass-specific metabolic rates (Lagos et al. 2017), and be more efficient in both energy conversion and food intake (Kooijman 2010; Taylor & Dunn 2018) with ecomorphological advantages for feeding efficiency (Luger et al. 2020). Whether these findings are congruent with our results needs further research on the link between metabolism and FR parameters.

Another important finding of our study concerns the negative consequences that climate change could have on recipient communities. The relationship between space clearance rate and temperature was indeed significantly negative in non-invasive species but tended to be positive in invasive species tested in their invasion range. This means that a warmer temperature could exacerbate the difference in FR and likely the asymmetry in competition between non-invasive (NIS) and invasive (IS-IR) species in recipient ecosystems. Our result brings support to the literature suggesting that invasive species would often be less vulnerable to warming because of predisposing physiological patterns, including energetic performance and thermal tolerance (Bates et al. 2013; Kelley 2014; Komoroske et al. 2021; Penk et al. 2019), their establishment being favored in certain ecosystemic contexts (Keller & Shea 2021; Sorte et al. 2010; Stachowicz et al. 2002).

To conclude, our study calls into question the shortcut that invasive species have higher FR than non-invasive species, without considering whether they were studied in the native or invasion range. Ecological filters leading to trait convergence (at least from the FR perspective) between invasive populations and resident non-invasive species seem to be at work in recipient communities. A promising research direction would be to compare species traits, especially metabolism, between donor and introduced populations of the same invasive species, and between recently introduced and long-established invasive populations. More broadly, this raises the question of how the invasion process shapes ecological niches through time (Ackerly 2003; Broennimann et al. 2007; Mukherjee et al. 2012; Stiels et al. 2015; Wiens & Graham 2005). Furthermore, the absence of differential FR between invasive species in the invasion range and non-invasive species raises concerns about the exclusive use of comparative FR to predict the trophic impact of an invasive species. FR should be used in combination with other metrics, such as species abundance, to better predict impact (Dickey et al. 2020; Laverty et al. 2017). Our study also reveals that global warming might exacerbate asymmetric competition between invasive and non-invasive species as space clearance rate was found to decrease with temperature in non-invasive species. We focused on freshwater fishes towards which FR literature is largely uneven and biased, probably because freshwater research offers ideal model species and experimental conditions for FR derivation. Extending our approach to other taxa is needed to know to what extent our results reflect general patterns.

## Author contributions

Marine A. Courtois and Chloé Souques equally contributed to this work. François-Xavier Dechaume-Moncharmont, Loïc Teulier and Vincent Médoc conceived the original idea and developed the protocol. Marine A. Courtois and Chloé Souques conducted bibliographical research and data collection. Marine A. Courtois and François-Xavier Dechaume-Moncharmont performed statistical analyses. Marine A. Courtois and Chloé Souques led the writing of the manuscript largely supported by François-Xavier Dechaume-Moncharmont, Loïc Teulier, Vincent Médoc and Yann Voituron. All authors analyzed the data, contributed critically to the MS and gave final approval for publication.

## Acknowledgements

This work was supported by the Heatwaves project and the BiodiverSâone project of the Agence de l’Eau Rhône-Méditerranée-Corse (AE-RMC). We thank Tristan Lefébure, Julien Clavel and Ian Dworkin for helpful advice on statistical analyses. We are grateful to the two anonymous reviewers for their careful evaluation and constructive feedback, which substantially strengthened this manuscript.

## Statement on inclusion

This study is based exclusively on publicly available datasets. As such, the geographic, taxonomic, and ecological representation of the data reflect the scope and limitations of the original data sources. We acknowledge that some regions and taxa may be under-represented and encourage future data collection efforts that broaden global and ecological coverage.

## Data availability statement

Data available from the Zenodo Repository: https://doi.org/10.5281/zenodo.18377688 (Courtois et al. 2026).

## SUPPLEMENTARY MATERIAL

**Fig. S1.**
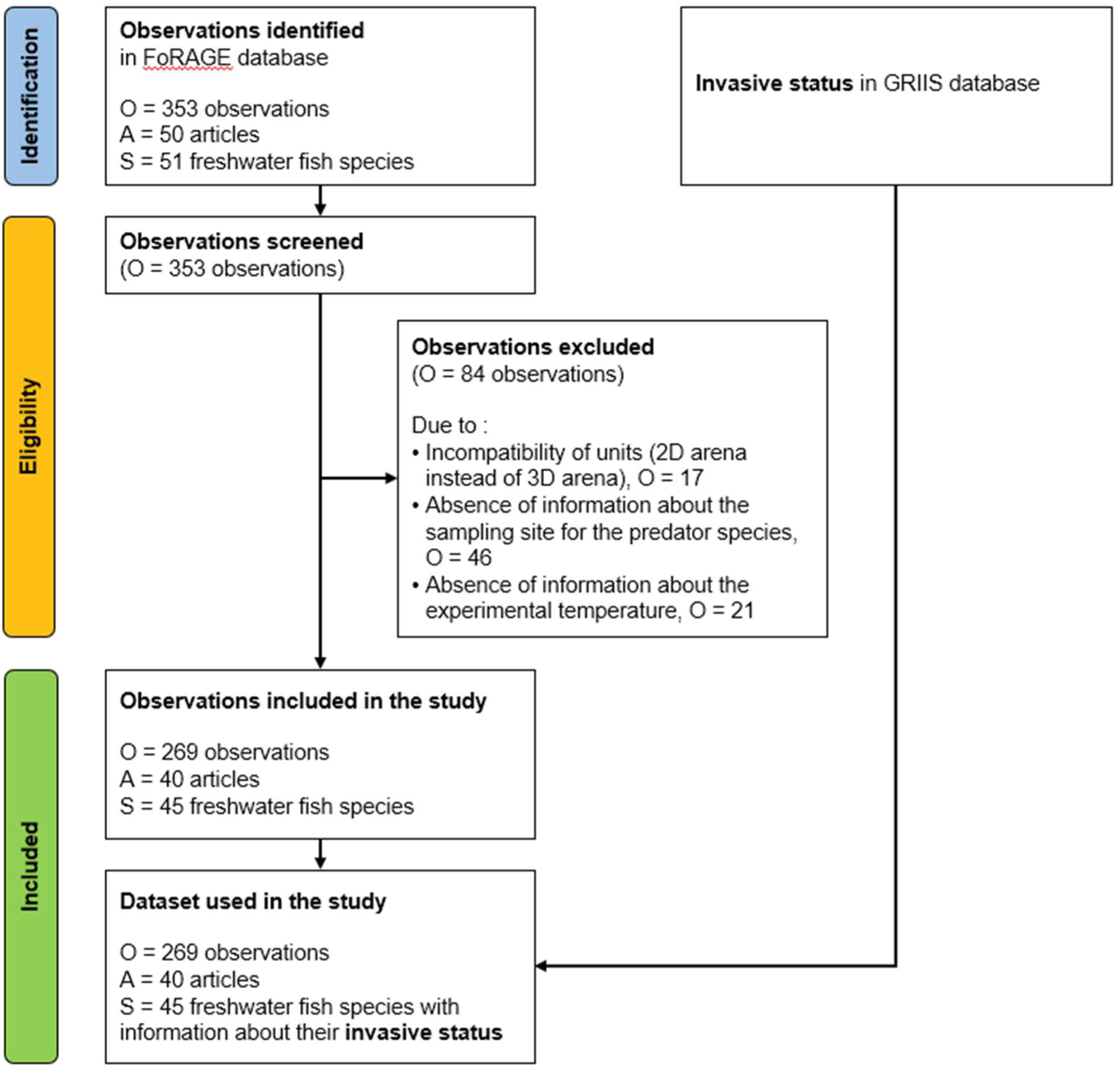
PRISMA flowchart showing the number of observations (O, i.e. number of estimated functional response curves reporting the space clearance rate a and the handling time h), the number of articles (A), and the number of freshwater fish species as predator (S).

**Figure S2.**
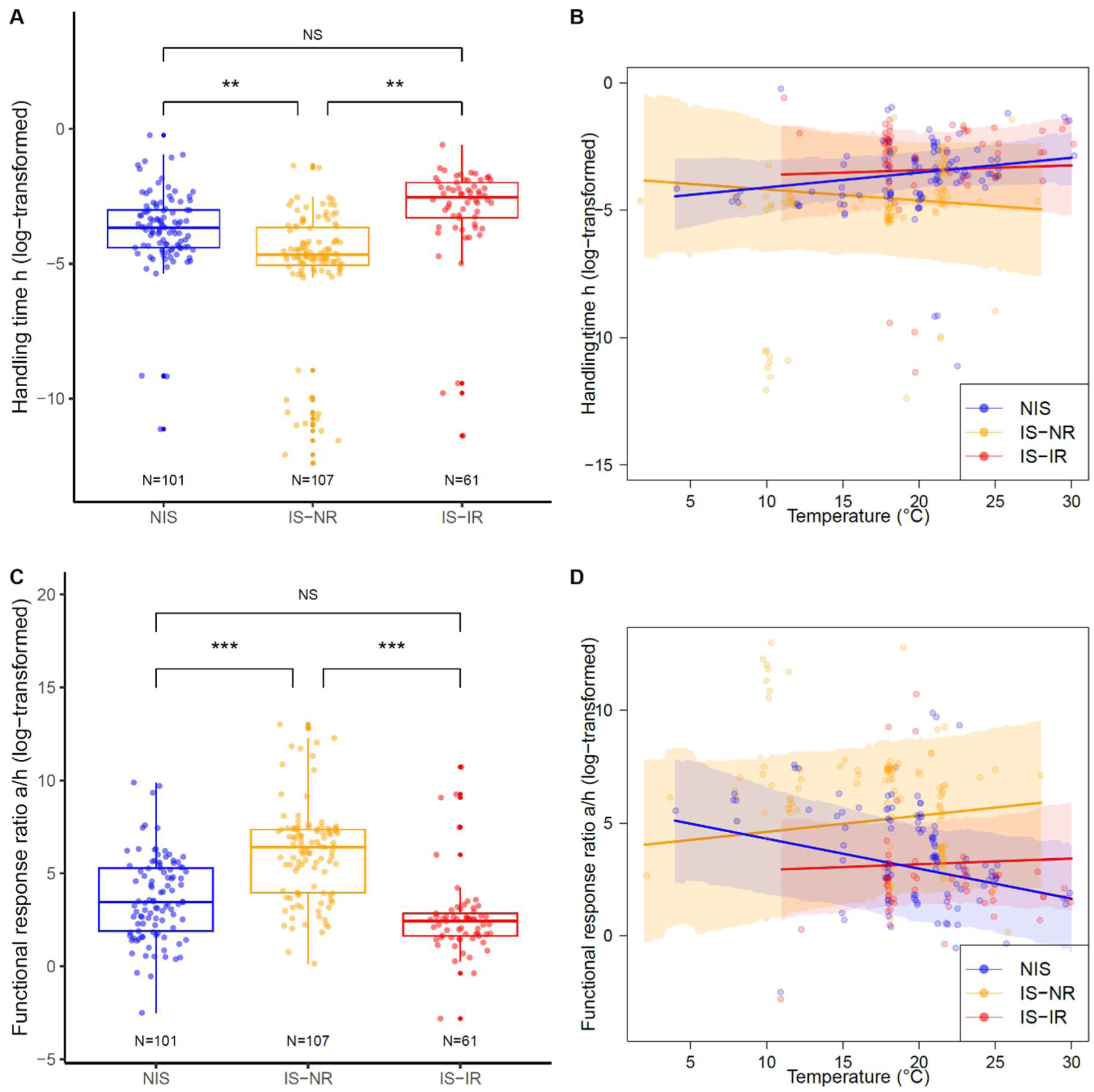
The functional response parameters are expressed in standard units (log-transformed). **A)** Boxplot of handling time (h) as a function of the sampling status, which was either non-invasive (NIS, in blue, all were sampled in their native range), invasive species sampled in their native range (IS-NR, donor populations, in orange) or invasive species sampled in their invasion range (IS-IR, introduced populations, in red). **B)** Effect of the interaction between sampling status and temperature on the handling time. **C)** Boxplot of functional response ratio (FRR = a/h) as a function of the sampling status. **D)** Effect of the interaction between sampling status and temperature on the functional response ratio (FRR = a/h). The sample size (N) is indicated below each group when relevant. Each data point corresponds to the report in a specific article of one estimated value of space clearance rate for one species. Multiple data points may correspond to the same species studied across different articles. The mixed-effect model incorporates multiple imputations and accounts for phylogenetic inertia. *** indicates pMCMC < 0.001, ** pMCMC < 0.01 and NS pMCMC > 0.05.

**Figure S3.**
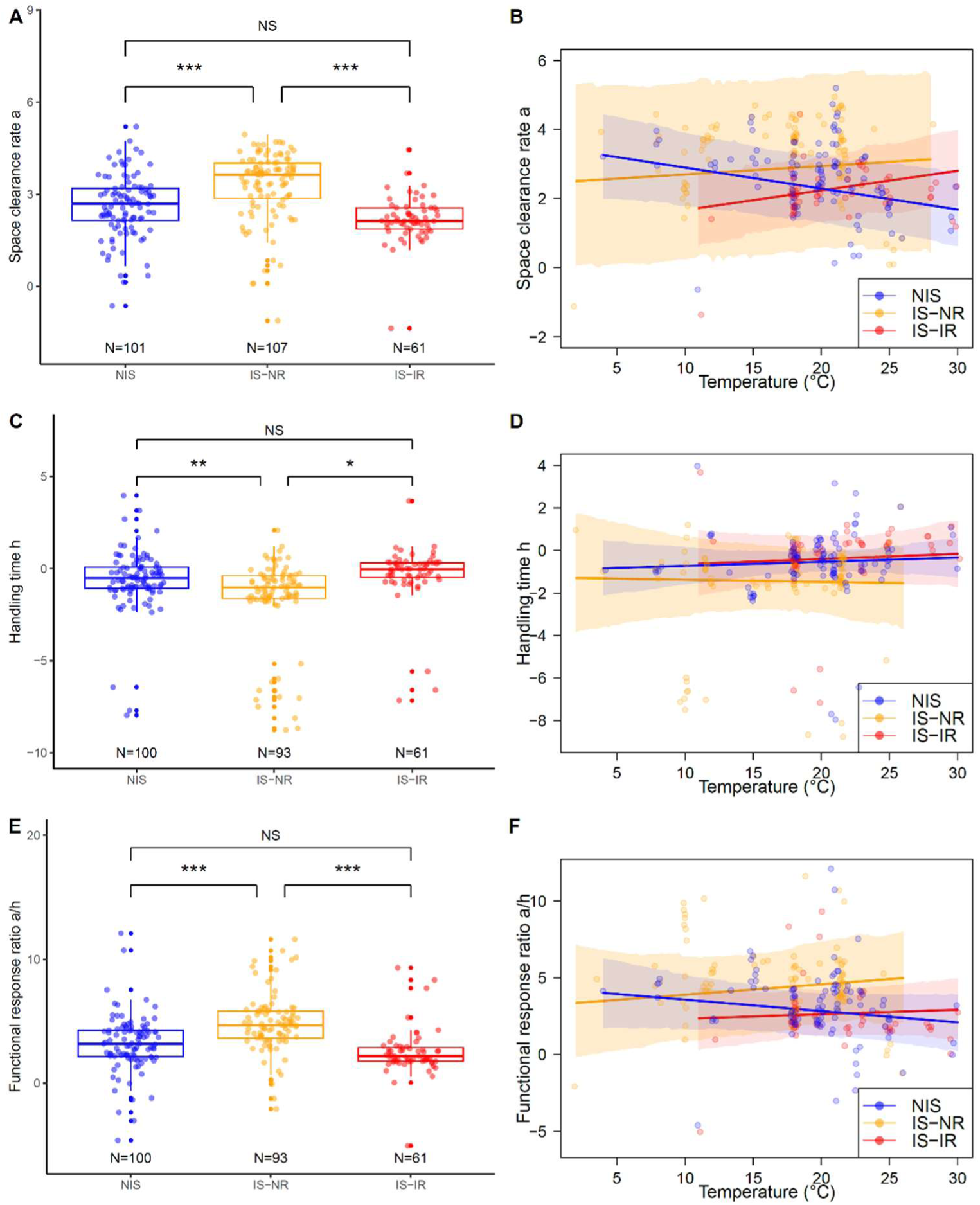
The functional response parameters are expressed in biomass units (log-transformed). **A)** Boxplot of space clearance rate as a function of the sampling status, which was either non-invasive (NIS, in blue, all were sampled in their native range), invasive species sampled in their native range (IS-NR, donor populations, in orange) or invasion species sampled in their invasive range (IS-IR, introduced populations, in red). **B)** Effect of the interaction between sampling status and temperature on the space clearance rate (a). **C)** Boxplot of handling time (h) as a function of the sampling status. **D)** Effect of the interaction between sampling status and temperature on the handling time (h). **E)** Boxplot of functional response ratio (FRR = a/h) as a function of the sampling status. **F)** Effect of the interaction between sampling status and temperature on the functional response ratio (FRR = a/h). The sample size (N) is indicated below each group when relevant. Each data point corresponds to the report in a specific article of one estimated value of clearance rate for one species. Multiple data points may correspond to the same species studied across different articles. The mixed-effect model incorporates multiple imputations and accounts for phylogenetic inertia. *** indicates pMCMC < 0.001, ** pMCMC < 0.01, * pMCMC < 0.05, and NS pMCMC > 0.05.

**Figure S4:**
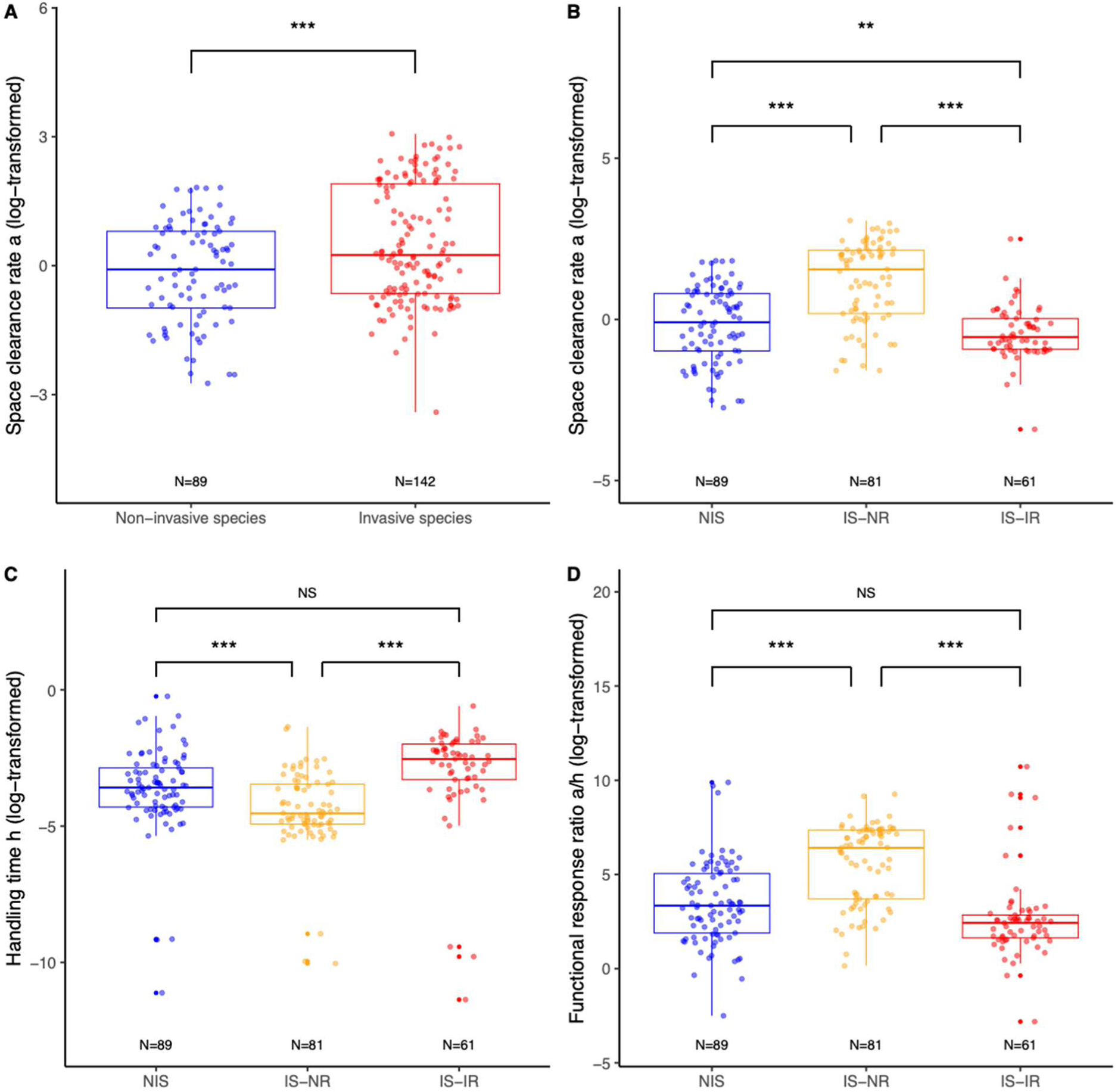
The functional response parameters are expressed in standard units (log-transformed). This figure corresponds to the analysis excluding Salmonidae. **A)** Boxplot of space clearance rate as a function of invasive status: non-invasive vs. invasive species. Invasive species had significantly higher functional response (FR) than non-invasive species, with a higher space clearance rate a (posterior mean = -1.15, 95% credible interval [95% CrI] = [-1.74; - 0.60], pMCMC = 3 × 10⁻⁴). **B–D)** Boxplots of space clearance rate a, handling time h, and functional response ratio a/h as a function of sampling status: non-invasive species (NIS, in blue, all sampled in their native range), invasive species sampled in their native range (IS-NR, donor populations, in orange), or invasive species sampled in their invasion range (IS-IR, introduced populations, in red). When invasive species were separated into donor (IS-NR) and introduced (IS-IR) populations, donor populations showed higher space clearance rate a (posterior mean = -1.17, 95% CrI = [-1.53; -0.76], pMCMC < 3×10⁻⁴ ***), lower handling time h (posterior mean = 1.06, 95% CrI = [0.47; 1.70], pMCMC = 3× 10⁻⁴), and higher functional response ratio a/h (posterior mean = -2.84, 95% CrI = [-3.68; -1.86], pMCMC < 3×10⁻⁴) than introduced populations. Introduced populations and native species do not differ significantly in space clearance rate a(posterior mean = -0.09, 95% CrI = [-0.60; -0.43], pMCMC = 0.71, NS), handling time h (posterior mean = -0.032, 95% CrI = [-0.73; 0.65], pMCMC = 0.91) or functional response ratio a/h (posterior mean = -0.69, 95% CrI = [-1.63; 0.31], pMCMC = 0.16). The sample size (N) is indicated below each group when relevant. Each data point corresponds to one estimated value of space clearance rate for a species reported in a specific article; multiple points may represent the same species across different studies. Mixed-effect models incorporate multiple imputations and account for phylogenetic inertia. *** indicates pMCMC < 0.001, and NS indicates pMCMC > 0.05.

**Figure S5:**
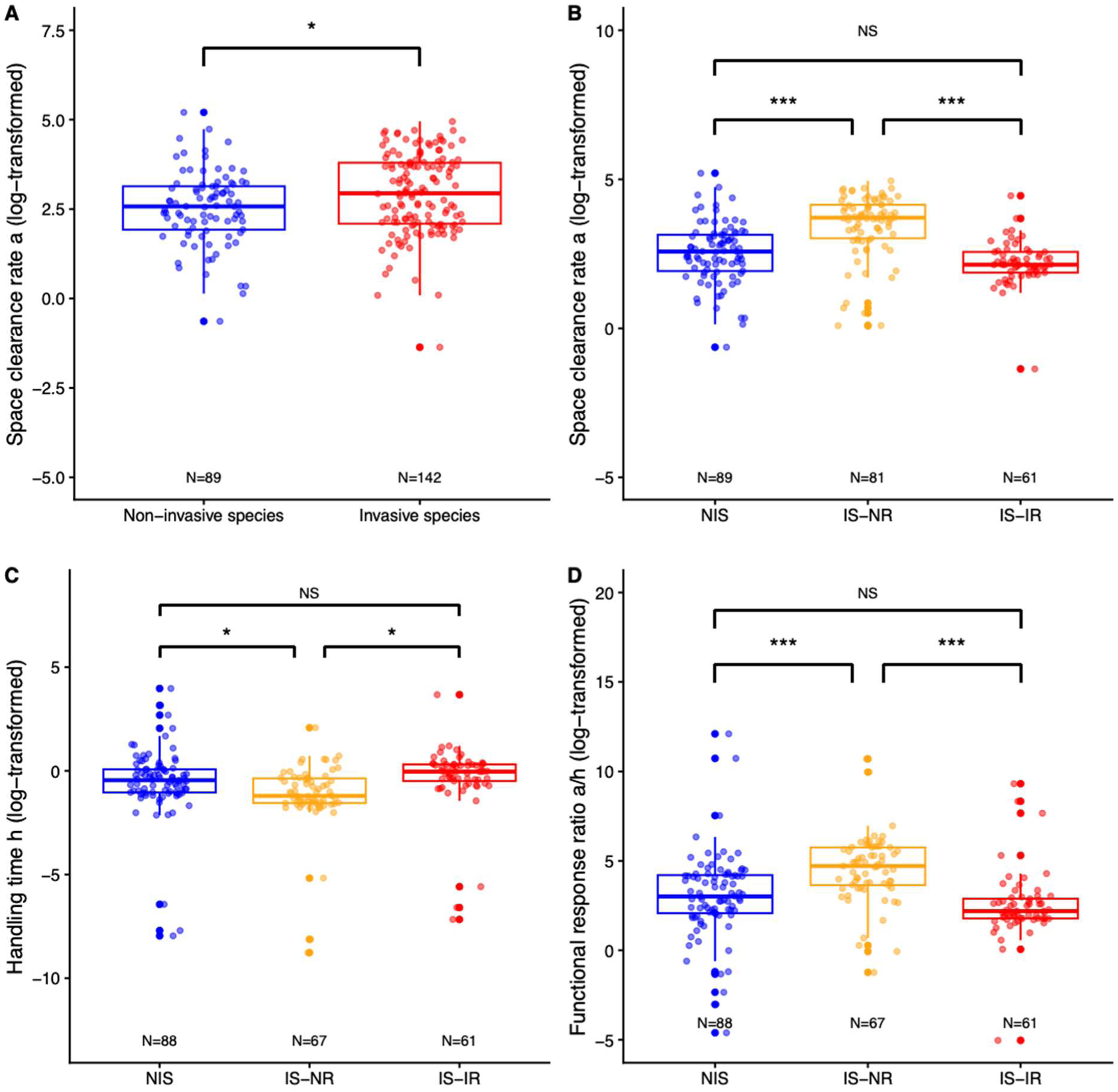
The functional response parameters are expressed in biomass units (log-transformed). This figure corresponds to the analysis excluding Salmonidae. **A)** Boxplot of space clearance rate as a function of invasive status: non-invasive vs. invasive species. Invasive species had significantly higher functional response (FR) than non-invasive species, with a higher space clearance rate a (posterior mean = -0.68, 95% credible interval [95% CrI] = [-1.21; - 0.22], pMCMC = 0.012). **B–D)** Boxplots of space clearance rate a, handling time h, and functional response ratio a/h as a function of sampling status: non-invasive species (NIS, in blue, all sampled in their native range), invasive species sampled in their native range (IS-NR, donor populations, in orange), or invasive species sampled in their invasion range (IS-IR, introduced populations, in red). When invasive species were separated into donor (IS-NR) and introduced (IS-IR) populations, donor populations showed higher space clearance rate a (posterior mean = -0.93, 95% CrI = [-1.31; -0.59], pMCMC < 3×10⁻⁴), lower handling time h (posterior mean = 0.80, 95% CrI = [0.20; 1.43], pMCMC = 0.015), and higher functional response ratio a/h (posterior mean = -1.90, 95% CrI = [-2.81; -1.06], pMCMC < 3×10⁻⁴) than introduced populations. Introduced populations and native species differ significantly in space clearance rate a (posterior mean = -0.74, 95% CrI = [-1.29; -0.20], pMCMC = 0.0060) but not in handling time h (posterior mean = -0.12, 95% CrI = [-0.75; 0.52], pMCMC = 0.73) and functional response ratio a/h (posterior mean = -0.14, 95% CrI = [-1.06; 0.91], pMCMC = 0.77). The sample size (N) is indicated below each group when relevant. Each data point corresponds to one estimated value of space clearance rate for a species reported in a specific article; multiple points may represent the same species across different studies. Mixed-effect models incorporate multiple imputations and account for phylogenetic inertia. *** indicates pMCMC < 0.001, and NS indicates pMCMC > 0.05.

**Figure S6.**
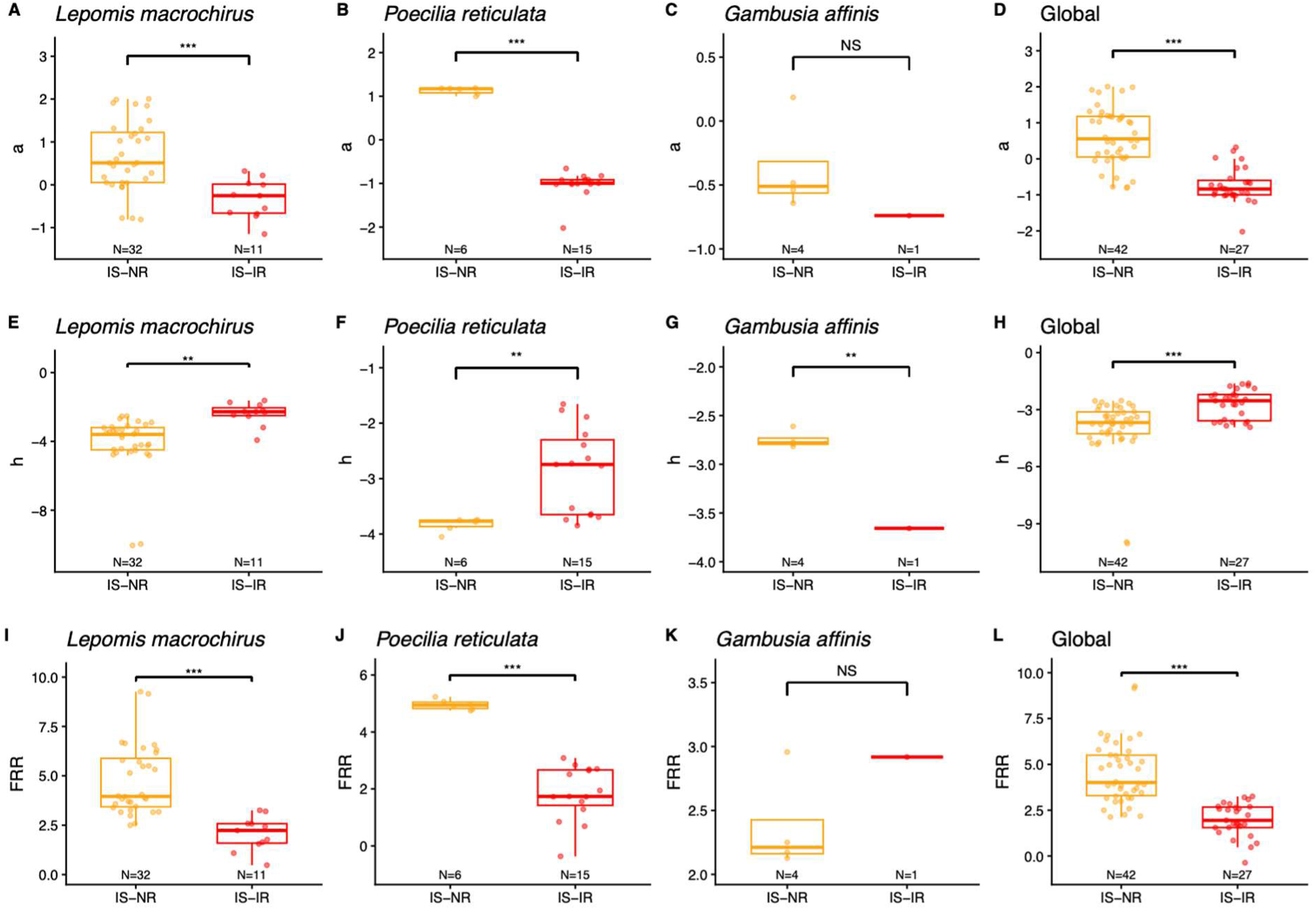
Comparison of functional response parameters (expressed in standard units) between native (IS-NR) and invasion (IS-IR) populations for the three species sampled in both ranges. The functional response parameters are expressed in standard units (log-transformed). Panels **A, B, C, D** show the space clearance rate **a**, panels **E, F, G, H** show the handling time **h**, and panels **I, J, K, L**show the functional response ratio **a/h**. Panels **A, E, I** correspond to Lepomis macrochirus, panels **B, F, J** to Poecilia reticulata, and panels **C, G, K** to Gambusia affinis. Panels **D, H, L** show a global analysis including all three species, with species included as a random factor. Points represent individual populations. Statistical significance for species-specific tests is indicated using p-values (*** indicates p < 0.001, ** p < 0.01, * p < 0.05, NS p > 0.05), whereas significance for the global model is indicated using pMCMC values.

**Figure S7.**
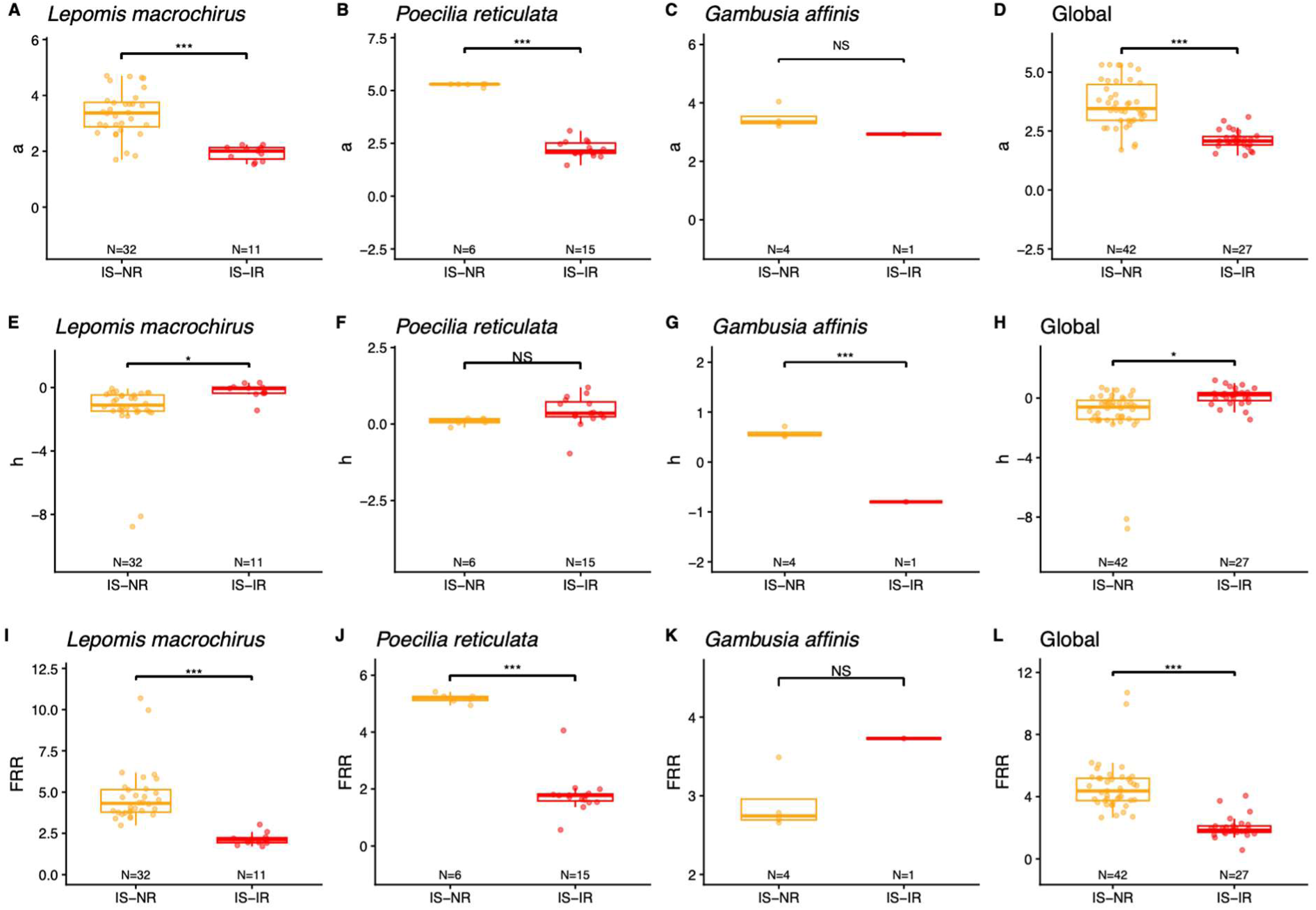
Comparison of functional response parameters (expressed in biomass units) between native (IS-NR) and invasion (IS-IR) populations for the three species sampled in both ranges. The functional response parameters are expressed in standard units (log-transformed). Panels **A, B, C, D** show the space clearance rate **a**, panels **E, F, G, H** show the handling time **h**, and panels **I, J, K, L**show the functional response ratio **a/h**. Panels **A, E, I** correspond to Lepomis macrochirus, panels **B, F, J** to Poecilia reticulata, and panels **C, G, K** to Gambusia affinis. Panels **D, H, L** show a global analysis including all three species, with species included as a random factor. Points represent individual populations. In the global analysis including the three species, with species included as a random factor and invasive populations separated into donor (IS-NR) and introduced (IS-IR) populations, donor populations showed higher space clearance rate a (posterior mean = 1.84, 95% CrI = [1.47; 2.26], pMCMC < 3×10⁻⁴), lower handling time h (posterior mean = -0.83, 95% CrI = [-1.61; -0.05], pMCMC =0.04), and higher FRR a/h (posterior mean = 2.60, 95% CrI = [1.92; 3.25], pMCMC < 3×10⁻⁴) than introduced populations. Statistical significance for species-specific tests is indicated using p-values (*** indicates p < 0.001, ** p < 0.01, * p < 0.05, NS p > 0.05), whereas significance for the global model is indicated using pMCMC values.

**Table S1.**
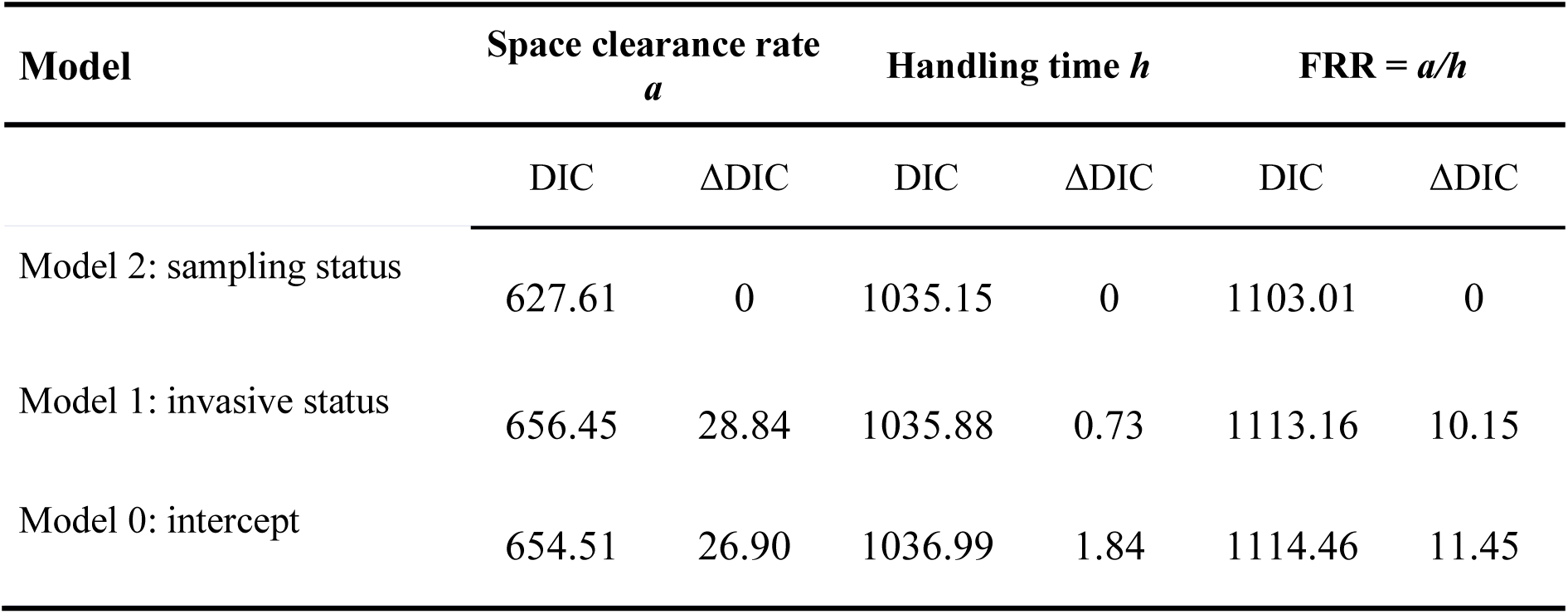
Model selection table for the influence of sampling status on each functional response parameter (space clearance rate a, handling time h, and functional response ratio FRR = a/h) expressed in biomass units. The competitive models were ranked according to their DIC. For a detailed description of the three models, refer to the main text.

| Model | Space clearance rate<br>$a$ | | Handling time $h$ | | FRR = $a/h$ | |
| --- | --- | --- | --- | --- | --- | --- |
| | DIC | ΔDIC | DIC | ΔDIC | DIC | ΔDIC |
| Model 2: sampling status | 627.61 | 0 | 1035.15 | 0 | 1103.01 | 0 |
| Model 1: invasive status | 656.45 | 28.84 | 1035.88 | 0.73 | 1113.16 | 10.15 |
| Model 0: intercept | 654.51 | 26.90 | 1036.99 | 1.84 | 1114.46 | 11.45 |

**Table S2.** Model selection table for the effect of sampling status and experimental temperature on each functional response parameter (space clearance rate a, handling time h, and functional response ratio FRR = a/h) expressed in biomass units. The full model (‘Model 3’, see main text) and all the nested models were ranked according to their DIC. The fixed variables considered were the experimental temperature (T°), the sampling status (NIS, IS-NR, IS-IR), and their interaction term (T°:status). Degree of freedom (df) for each model is also reported.

| Parameter | Nested model | | | df | DIC | ΔDIC |
| --- | --- | --- | --- | --- | --- | --- |
| $a$ | $T^\circ$ | Status | $T^\circ:Status$ | 8 | 609.7 | 0 |
|  |  | Status |  | 5 | 627.5 | 17.85 |
| | $T^\circ$ | Status | | 6 | 629.3 | 19.64 |
|  |  |  |  | 3 | 654.6 | 44.94 |
| | $T^\circ$ | | | 4 | 656.1 | 46.47 |
| $h$ | | Status | | 5 | 1035.2 | 0 |
|  |  |  |  | 3 | 1037.0 | 1.83 |
| | $T^\circ$ | Status | | 6 | 1038.0 | 2.81 |
| | $T^\circ$ | | | 4 | 1040.2 | 5.06 |
| | $T^\circ$ | Status | $T^\circ:Status$ | 8 | 1040.9 | 5.72 |
| FRR $a/h$ | $T^\circ$ | Status | $T^\circ:Status$ | 8 | 1100.8 | 0 |
|  |  | Status |  | 5 | 1103.0 | 2.21 |
| | $T^\circ$ | Status | | 6 | 1104.9 | 4.60 |
|  |  |  |  | 3 | 1114.4 | 13.58 |
| | $T^\circ$ | | | 4 | 1116.7 | 15.87 |

**Table S3.** Effect of the experimental temperature for each functional response parameter (space clearance rate a, handling time h, and functional response ratio FRR = a/h) expressed in standard units, and for each sampling status (NIS, IS-NR, IS-IR). The posterior mean of the slope with 95% credible interval, and pMCMC are reported. *** indicates pMCMC < 0.001, ** pMCMC < 0.01, * pMCMC < 0.05.

| Parameter | Sampling status | Posterior mean | 95% CrI | pMCMC |
| --- | --- | --- | --- | --- |
| $a$ | NIS | -0.095 | [-0.14, -0.052] | $< 3 \times 10^{-4}$ *** |
|  | IS-NR | 0.0078 | [-0.046, 0.056] | 0.767 |
|  | IS-IR | 0.056 | [-0.0073, 0.11] | 0.072 |
| $h$ | NIS | 0.054 | [-0.019, 0.13] | 0.15 |
|  | IS-NR | -0.044 | [-0.19, 0.087] | 0.52 |
|  | IS-IR | 0.020 | [-0.11, 0.16] | 0.76 |
| FRR $a/h$ | NIS | -0.13 | [-0.11, -0.011] | 0.014 * |
|  | IS-NR | 0.069 | [-0.062, 0.20] | 0.32 |
|  | IS-IR | 0.025 | [-0.12, 0.18] | 0.75 |

**Table S4.** Effect of the experimental temperature for each functional response parameter (space clearance rate a, handling time h, and functional response ratio FRR = a/h) expressed in biomass units, and for each sampling status (NIS, IS-NR, IS-IR). The posterior mean of the slope with 95% credible interval, and pMCMC are reported. * indicates pMCMC < 0.05.

| Parameter | Sampling status | Posterior mean | 95% CrI | pMCMC |
| --- | --- | --- | --- | --- |
| $a$ | NIS | -0.060 | [-0.11, -0.011] | 0.017 * |
|  | IS-NR | 0.025 | [-0.029, 0.075] | 0.34 |
|  | IS-IR | 0.056 | [-0.0034, 0.12] | 0.072 |
| $h$ | NIS | 0.018 | [-0.061, 0.090] | 0.61 |
|  | IS-NR | -0.010 | [-0.17, 0.14] | 0.96 |
|  | IS-IR | 0.024 | [-0.087, 0.14] | 0.68 |
| FRR $a/h$ | NIS | -0.074 | [-0.19, 0.035] | 0.20 |
|  | IS-NR | 0.067 | [-0.12, 0.24] | 0.51 |
|  | IS-IR | 0.030 | [-0.11, 0.19] | 0.69 |

